# Islet size, cellular composition, and spatial location are interconnected in the human pancreas

**DOI:** 10.64898/2026.10.01.756003

**Authors:** Sean D. Dufault-Hunter, Vipul Periwal, Manami Hara

## Abstract

Stem cell-derived islets (SC-islets), a promising therapy for type-1 diabetes, are often dissociated and reaggregated into standardized sizes. However, the relationship between endocrine cell composition and size has not been precisely defined in native human islets. In this study, we examined relationships between islet size, composition, and spatial distribution throughout the whole pancreata of 10 well-characterized and biometrically diverse donors. Insulin, glucagon, and somatostatin stains were used to identify the major endocrine cell types. Islets ranging in size from non-islet single cells and small clusters (niSSCs) to >500 μm were analyzed. Islet composition strongly correlated with islet size, with large islets having increased alpha-cell fractions. Delta-cell fraction was consistent across sizes. Approximately 80% of the total endocrine area was contained in islets positive for all three hormones. Spatial analysis additionally revealed clustering among niSSCs, tri-hormone-positive islets, and cells/clusters composed of a single cell type.

## Introduction

SC-islets have emerged as a promising cell replacement therapy for T1D, prompting intensive efforts to engineer therapeutic grafts that closely recapitulate the architecture and function of native human islets (1–4). To improve reproducibility and transplantation outcomes, SC-islet organoids are frequently dissociated and reaggregated into uniformly sized clusters (1, 5, 6). In parallel, considerable efforts have been directed toward optimizing endocrine cell composition, particularly the relative abundance of beta- and alpha-cells, to enhance graft function (7, 8). These strategies implicitly assume that islet size and cellular composition are key determinants of endocrine function. Prior literature suggests that human islets exhibit size-dependent differences in islet cell composition (9–13). However, while it has been recognized, this fundamental morphological relationship has yet to be precisely codified for all three major endocrine cell types. Defining this association is critical for establishing rational design criteria for SC-islets, and to determine whether current engineering strategies reproduce the organization of native human islets.

Concurrently, substantial advances in the study of intraislet cell-cell communication have revealed a highly coordinated network of paracrine and contact-dependent interactions through which islet cell populations continuously modulate one another to preserve glucose homeostasis (14–17). Particularly, alpha-cell to beta-cell communication has emerged as a rapidly evolving area of research, with accumulating evidence indicating that alpha-cells actively support beta-cell function, especially through glucagon and locally produced GLP-1-related signaling (18–20). Similarly, delta-cells are increasingly recognized as active regulators of islet glucose homeostasis rather than simply passive inhibitors of insulin and glucagon secretion (21). This reciprocal beta-cell-delta-cell feedback involves beta-cell-derived urocortin-3, which stimulates somatostatin release, allowing delta-cells to dynamically modulate beta-cell insulin secretion (22, 23).

A detailed understanding of the non-diseased organ is essential to specify the changes to islet morphology and overall pancreatic architecture that occur in disease states, including diabetes. Recognition of this fact has led groups to investigate more comprehensive datasets, even going as far as performing full 3D scans of pancreata (11, 24). In the present study, we use a well-characterized, previously published dataset of 10 age-matched, male, non-diabetic humans (10, 25). Imaging was performed along the full length of the pancreas stereologically, allowing us to examine islets across the entire tissue. Further, these subjects vary greatly in BMI, age, and total endocrine cell mass. This diversity allows us to identify and specifically define relationships that are robust enough to persist across these biometric differences.

We applied bioinformatic approaches to characterize approximately 20,000 islets per subject, incorporating comprehensive information at the individual-islet level, which includes size (normalized to an effective diameter), cellular composition based on the expression of three major islet hormones, spatial X-Y coordinates within individual tissue sections (∼27 per subject), and anatomical location along the pancreatic head-body-tail axis. Spatial organization was further assessed using Ripley’s K, the empty-space function, and average border distance. Islets were classified according to their hormonal expression profiles. Positive signals for insulin (INS), glucagon (GCG), and somatostatin (SST) were used to form groups of mono-, di- and tri-hormonal islets, yielding seven distinct islet categories. The analyses also incorporated niSSCs, with particular attention to insulin-positive cells, which have recently been identified as the most vulnerable cell populations to autoimmune attack in T1D (12, 13).

## Results

### The majority of INS, GCG, and SST areas are found in tri-hormone-positive islets 100-200 μm in diameter

Large-scale imaging data was used to build a dataset containing size and compositional information on islets from 10 whole, non-diabetic human pancreata (donor information provided in Supplementary Table 1). Islets were organized into groups based on anatomical region, diameter and cellular composition. The proportions of total INS, GCG, and SST areas contained in each of these groups were quantified and visualized. First, data were divided into a series of bins using islet effect diameter (IED; normalized to a circle) cutoffs (40 μm, 100 μm, and 200 μm), and hormone areas in each bin were measured. The proportion of total area in each bin increased before reaching a peak in islets 100 – 200 μm in diameter (INS 40.1 ± 3.8%, GCG 48.0 ± 5.0%, and SST 42.4 ± 4.2%) and decreasing in islets > 200 μm in diameter (Figure 1A; Supplementary Table 2). More of the total GCG area appeared to be in islets > 100 μm in diameter as compared to the other two hormones (69.7 ± 10.1% of GCG area, 52.5 ± 6.8% of INS area, 58.1 ± 6.2% of SST area) (Figure 1A, Supplementary Table 2). Consistent with previous reports (11, 26), niSSCs (IED < 40 µm) accounted for 8.3 ± 1.8% of the total endocrine area despite them making up 47.6 ± 4.9% of all islets in the pancreas (Figure 1A; Supplementary Table 2).

**Figure 1:**
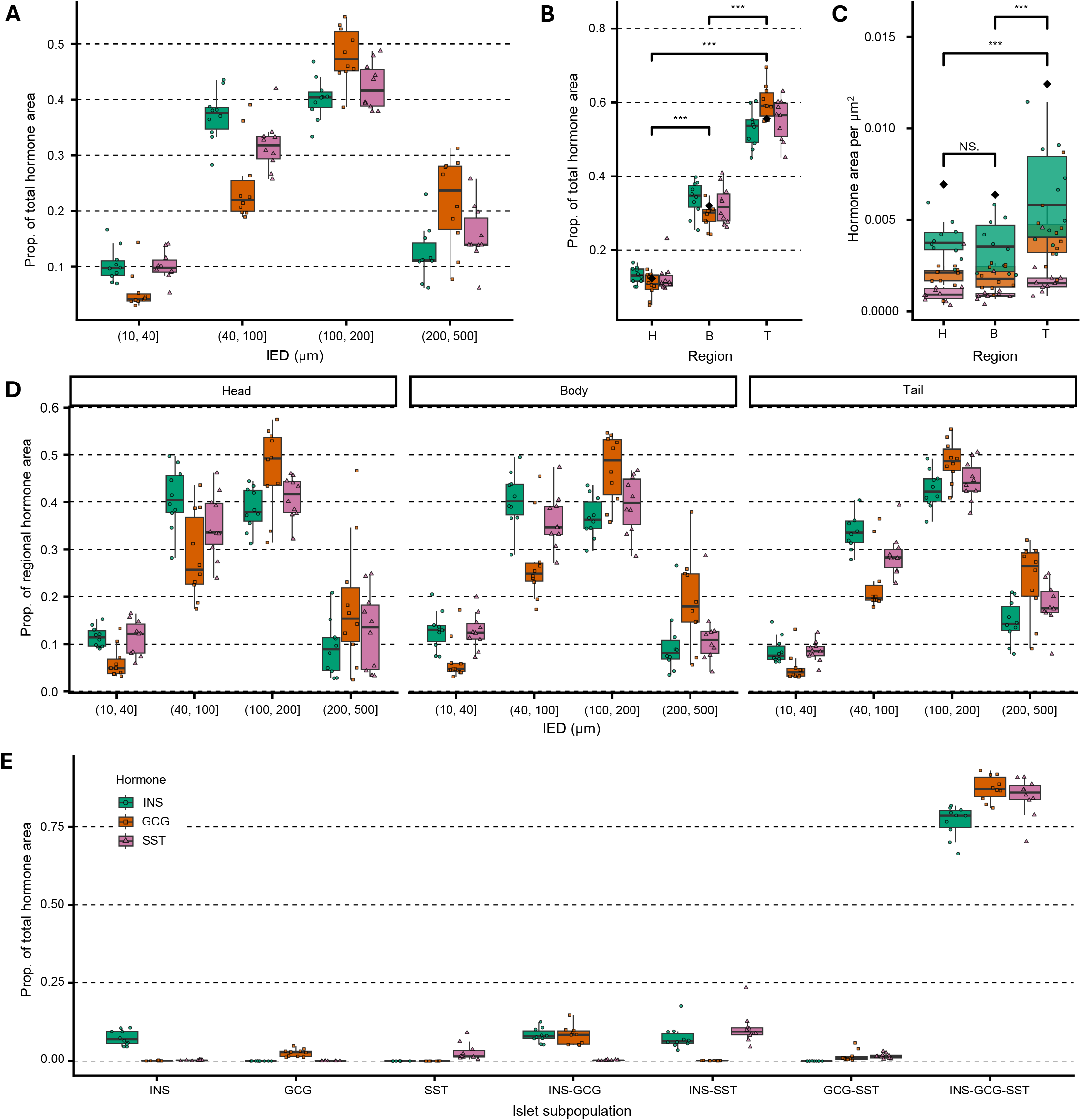
Relationship between proportion of INS, GCG, and SST areas and islet size. (**A**) Proportion of total hormone area in IED bins. Points on box-plots represent the mean value calculated from each subject. (**B**) Proportion of total hormone area in each region. Black diamonds represent mean overall hormone proportion. Significance of the differences between regions, as calculated by one-way ANOVAs, is presented. (**C**) Hormone area per µm^2^ in each region. Black diamonds represent mean overall hormone density. Significance of differences between regions, as calculated by one-way ANOVAs, is presented. (**D**) Proportion of regional hormone area in IED bins separated by region. (**E**) Proportion of total hormone area in each subpopulation.

Next, the proportions of each hormone in the head, body, and tail regions were examined. Significantly different proportions of total endocrine areas were contained in each region, increasing from the head and body to the tail region for all three major hormones (p < 0.001) (Figure 1B, Supplementary Table 2). In the tail, the proportion of GCG area (60.1 ± 4.6%) was greater than that of INS area (52.9 ± 5.1%) or SST area (55.2 ± 5.7%) (Figure 1B; Supplementary Table 2). The density of total hormone-positive area was approximately 2-fold higher in the tail region (Figure 1C, p < 0.001).

Diameter and anatomical region were also combined to determine whether the proportion of endocrine area in each bin differed between regions. Despite the substantial differences in total endocrine area, the relationship between hormone-positive area and islet diameter was conserved across the three pancreatic regions (Figure 1D).

A coarse-grained characterization of cellular composition was obtained by dividing up islets based on positivity for the hormone of each major cell type, resulting in seven subpopulations: INS, GCG, SST, INS-GCG, INS-SST, GCG-SST, and INS-GCG-SST. The majority of INS, GCG, and SST areas were contained within tri-hormone (INS-GCG-SST) islets (81.5 ± 4.5% of total endocrine area) (Figure 1E). The concentration of such a large proportion of endocrine cell mass within tri-hormone islets is unexpected, given the heterogeneity observed among other islet profiles (25, 27, 28).

### Distributions of islet size and hormone positivity are conserved across regions

Next, we sought to define the distributions of islet size and cellular composition in the full pancreas and in each region. Islets were again organized into groups, and the numerical frequency of each group was quantified and visualized. Frequency plots of diameter bins in the head, body, and tail regions revealed similar distributions, though there were more islets with diameters > 60 μm in the tail region (Figure 2A). Summary distributions for the whole pancreas were also computed for completeness (Supplemental Figure 1A-B). These plots agreed with the per-region data. The most numerically abundant size range was 21–30 μm (Figure 2A, Supplemental Figure 1A-B). This spike was followed by a downward slope as IED increased (Figure 2A, Supplemental Figure 1A-B). We observed a median diameter of 41.1 ± 25.6 μm across all islets in the dataset.

**Figure 2:**
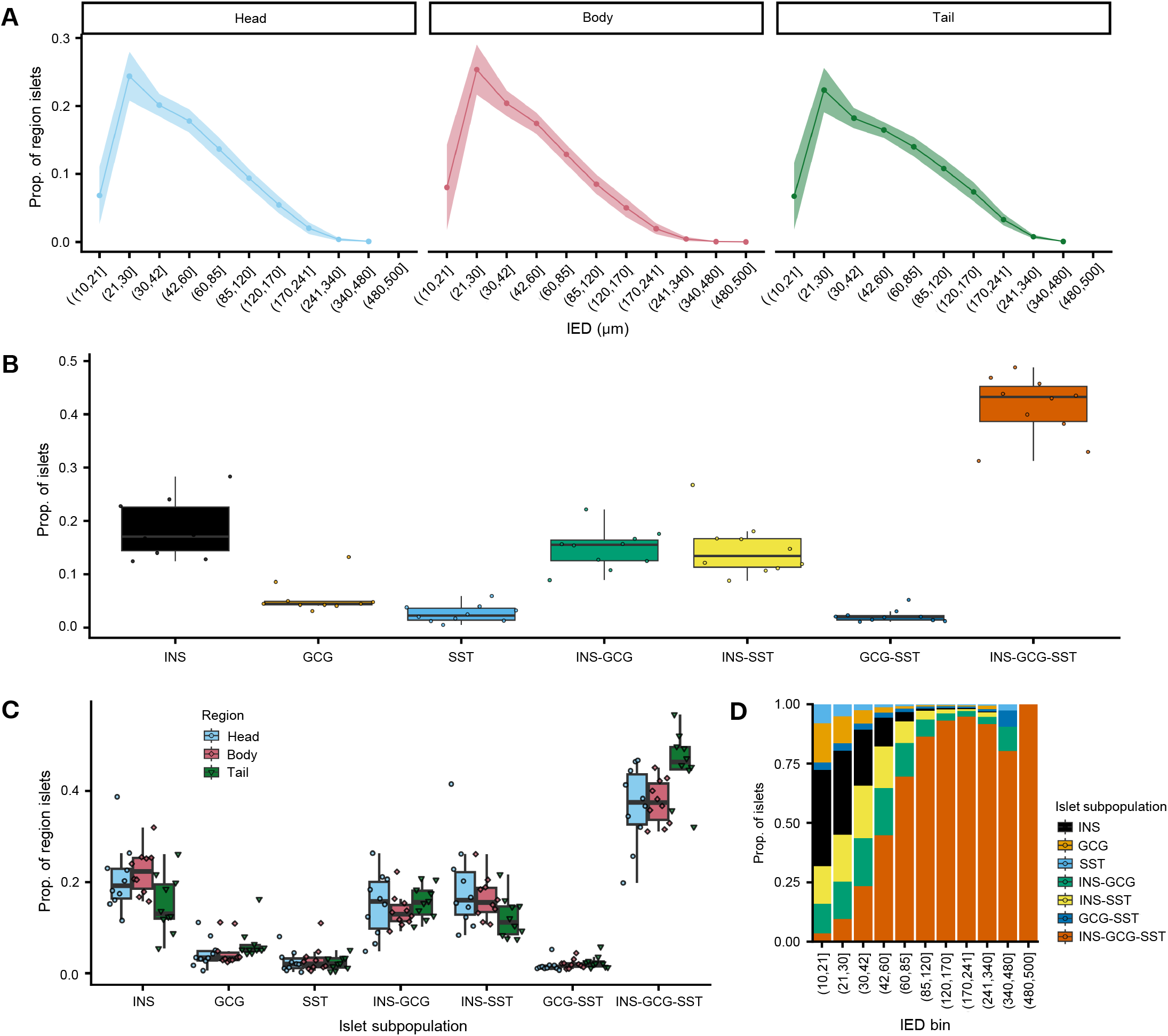
Distributions of islet subpopulations across regions. (**A**) Distributions of IED within each region. Points represent the mean value across all subjects; ribbons represent one standard deviation. (**B**) Proportion of each hormone-positivity-based subpopulation across donors. Points on box-plots represent the mean value computed from each donor. (**C**) Proportion of each hormone-based subpopulation, separated by region. Shapes and colors correspond to region. (**D**) Proportion of each hormone-based subpopulation within each IED bin (see A). Bars represent the mean between all 10 subjects.

Examination of the frequency distributions of each hormone-based subpopulation revealed that tri-hormone islets were the most common overall, with a mean frequency of 41.4 ± 5.8% across the ten subjects (Figure 2B). The most abundant of the mono-hormone subpopulations were INS-only islets at a mean frequency of 18.6 ± 5.4%, as compared to 5.6 ± 3.0% and 2.6 ± 1.6% for GCG- and SST-only islets respectively (Figure 2B, Supplementary Table 1). These patterns were broadly consistent across anatomical regions, although the tail region had a greater proportion of tri-hormone islets than the head and body (Figure 2C).

Following these analyses, the relationship between islet size and cellular composition was examined by quantifying the frequency of each hormone-based subpopulation within each IED bin. Increasing islet size was associated with a marked shift in population composition. The frequency of mono- and di-hormone islets decreased with increasing islet size whereas tri-hormone islets became predominant, rising from 12.8 ± 4.7% in the smallest bin to 97.2 ± 2.4% in the largest (Figure 2D, Supplementary Table 2). Smaller bins (10 – 85 μm) were more diverse. INS-GCG and INS-SST islets had roughly equal frequency among these bins (Figure 2D, Supplementary Table 2), while GCG-SST islets were rare. Of the mono-hormone islets, the INS subpopulation was the most prevalent.

### Beta- and alpha-cell proportions intersect in larger islets while delta-cell proportion is constant

Defining the relationships between cellular composition, islet diameter, and anatomical region required a finer survey of islet cytoarchitecture. We therefore examined the fraction of each islet’s area occupied by INS, GCG, and SST, providing approximate per-islet beta-, alpha-, and delta-cell ratios. The average fractions of each major cell type were first quantified in each diameter bin. This revealed that mean per-islet GCG fraction increased with diameter, whereas mean INS fraction decreased (Figure 3A; Supplementary Table 3). In some subjects, the mean GCG fraction even matched or surpassed the mean INS fraction in islets approximately 200 μm in diameter (Figure 3A). In these cases, mean fractions of both cell types were approximately 40–45% (Supplementary Table 3). These shifts are in agreement with previous reports that larger islets contain a greater proportion of alpha-cells (9, 11), and support findings for the importance of alpha-cells to glucose regulation and pathology (29–33). In contrast to INS and GCG, mean SST fraction was similar across islet size bins, ranging from 18.1 ± 3.4% in the smallest bin to 17.6 ± 12.7% in the largest (Figure 3A; Supplementary Table 3).

**Figure 3:**
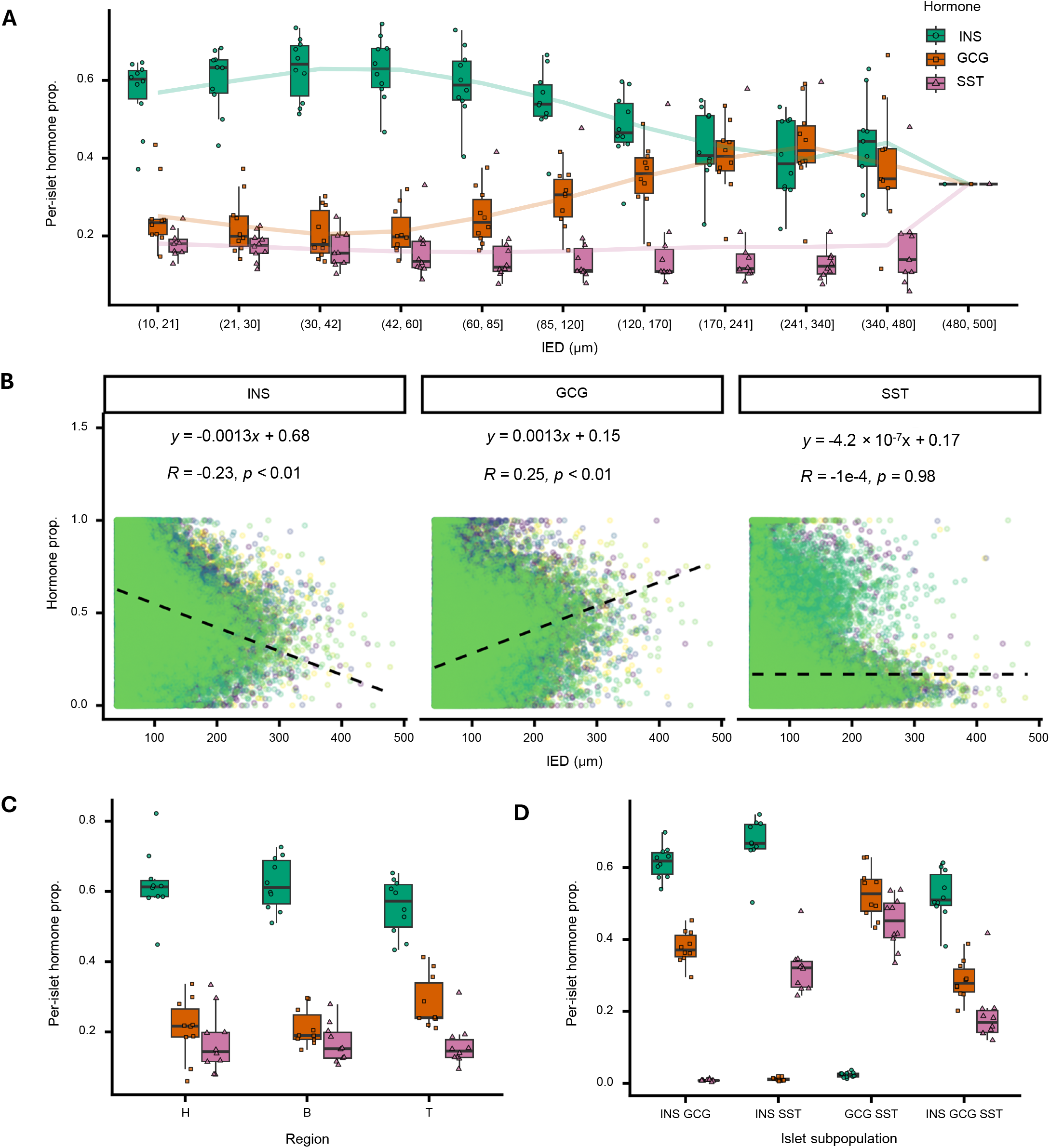
Relationship between cellular composition and islet size. (**A**) Average hormone fraction of each islet across diameter bins. Points on box-plots represent the mean value calculated from each subject. Lines trace the mean values across all subjects. (**B**) Linear regression with IED used as a predictor for hormone fraction. Presented equations exclude niSSCs. (**C**) Average hormone composition of an islet in each region. (**D**) Average hormone fraction of each islet in each di/tri-hormone islet subpopulation.

A linear regression using diameter as a predictor for INS, GCG, and SST fractions was performed to specifically quantify these relationships. INS fraction was negatively correlated with IED (slope = −0.0013, p < 0.01) and GCG fraction was positively correlated with IED (slope = 0.0013, p < 0.01) (Figure 3B). SST fraction did not appear to correlate with IED (slope = −4.2 × 10^-7^, p = 0.98) (Figure 3B). Islets < 100 μm in effective diameter showed greater variability in these fractions (Figure 3B). Consistent with the inter-subject variability observed throughout this study, the coefficients of determination revealed only modest correlation between hormone fractions and diameter (R² = 0.0529 for INS, 0.0625 for GCG, and 3.35 × 10^-4^ for SST). The expected composition of islets at a series of diameter values frequently used in SC-islet generation were computed using the fit models. Islets: (*i*) 100 μm in diameter had an expected composition of 54.1 ± 0.1% beta-cells, 28.6 ± 0.1% alpha-cells, and 17.3 ± 0.1% delta-cells, (*ii*) 150 μm in diameter had an expected composition of 48.6 ± 0.1% beta-cells, 34.6 ± 0.1% alpha-cells, and 16.8 ± 0.1% delta-cells, (*iii*) 200 μm in diameter had an expected composition of 43.2 ± 0.2% beta-cells, 40.6 ± 0.2% alpha-cells, and 16.2 ± 0.2% delta-cells, and (*iv*) 250 μm in diameter had an expected composition of 37.7 ± 0.3% beta-cells, 46.6 ± 0.3% alpha-cells, and 15.7 ± 0.2% delta-cells. We observed one notable outlier to the above trends. Performing the above analysis with data segmented by subject, rather than with all data pooled, showed that SST fraction rose sharply with effective diameter to overtake both INS and GCG fractions in donor 7 (Supplementary Figure 2).

Next, the mean fractional composition of islets in each region was examined. Average cell ratios appeared similar across anatomical location, with comparable mean INS, GCG, and SST fractions in islets from the pancreatic head, body, and tail (Figure 3C). There do not appear to be substantial compositional differences between islets drawn from different regions.

Average cell ratios were also quantified for the coarse approximations of islet composition. Plotting the average cell fractions of these groups revealed that beta-cells were the most abundant cell type in all INS subpopulations (Figure 3D). While this initially seems to contradict the findings presented in Figure 3A, a deeper examination reveals that they are in alignment. Islets < 100 μm in diameter make up a numerical majority of islets in the pancreas and are associated with a high INS fraction. Therefore, mean ratios in each of these subpopulations are skewed towards values associated with these bins; i.e. a high INS fraction.

### niSSCs, tri-hormone and mono-hormone islets cluster together

Beyond composition, we also sought to examine how islets are spatially distributed in the tissue. Islet coordinates were used to create representative point patterns for each tissue section, with metadata on diameter and hormone positivity appended to each point (islet) (Supplementary Figure 3A-B). Organizing the data in this way allowed for the application of statistics to estimate the spatial character of islet subpopulations.

Namely, Ripley’s K (H(r)) and the empty space function (F(x)) were used to estimate the degree of clustering/dispersion among points in these representative patterns. Both Ripley’s K and the empty-space function quantify clustering/dispersion at different scales. The output of these measures is therefore a function, interpreted using a confidence interval (p = 0.01) generated via a series of simulations. H(r) values above this threshold indicate clustering while values below indicate regular dispersion. The F(x) function has an opposite interpretation, with high values indicating regular dispersion and low values indicating clustering. A visual model of the mechanism behind these computations is provided in Supplementary Figure 3C.

Estimates of both functions varied across sections and subjects. However, review of the full set of more than 200 plots revealed consistent relationships between islet composition, size, and spatial organization. The most striking finding was the absence of evidence for regular dispersion. No significant (p = 0.01) estimate from any population supported regular dispersion. In contrast, most sections had evidence of clustering among one or more subpopulations (Table 1). Estimates derived from niSSCs (Figure 4A) and islets with effective diameters of 40 – 100 µm (Figure 4B) consistently showed significant evidence of clustering, meeting the threshold (p = 0.01) in 77.8% and 71.9% of sections respectively (Table 1). H(r) plots generated from these subpopulations followed a loose, shallow logarithmic curve and exceeded the significance threshold from approximately 250 µm onward (Figure 4A-B). F(x) plots showed evidence of clustering around the same value and resembled a fishhook, curving inwards towards zero after a dip.

**Figure 4:**
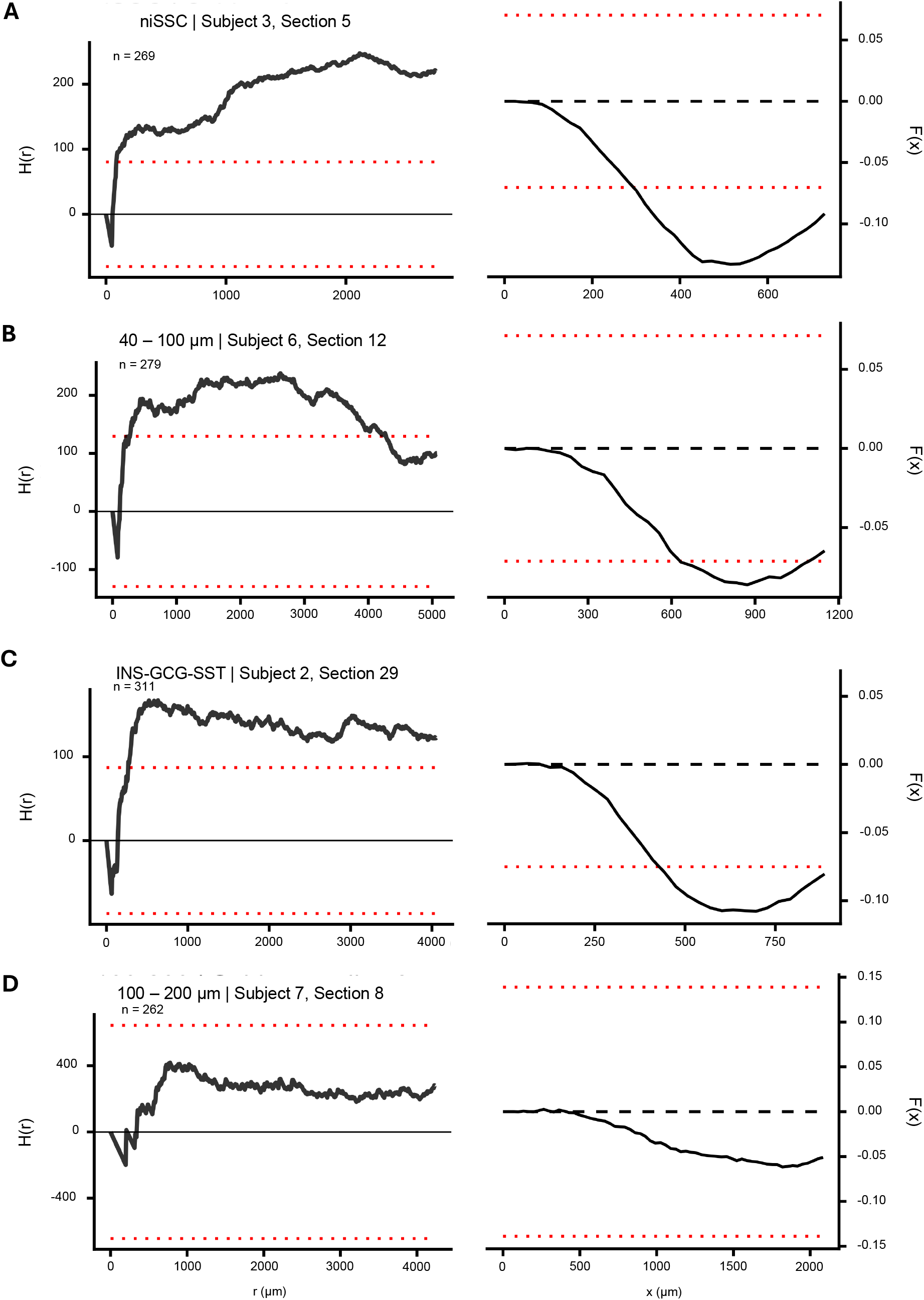
Spatial analysis of niSSCs, small, medium, large, and tri-hormone islets. (**A**) Representative spatial plots estimated using niSSCs. Ripley’s K simplified to the H function is presented on the left and the empty space function F(x) is presented on the right. The expected value under CSR for both functions is 0. The red dotted lines show a global confidence envelope generated from 99 simulations. Interpretations of the plots are reversed: An H(r) value above what is expected under CSR indicates clustering, while an F(x) value below what is expected under CSR indicates clustering. Islet count is shown. Subject and section are shown in the plot title (Subject number, Section number). (**B**) Representative spatial plots estimated using islets 40-100 μm in effective diameter. (**C**) Representative spatial plots estimated using tri-hormone islets. (**D**) Representative spatial plots estimated using islets 100-200 μm in effective diameter.

**Table 1:**
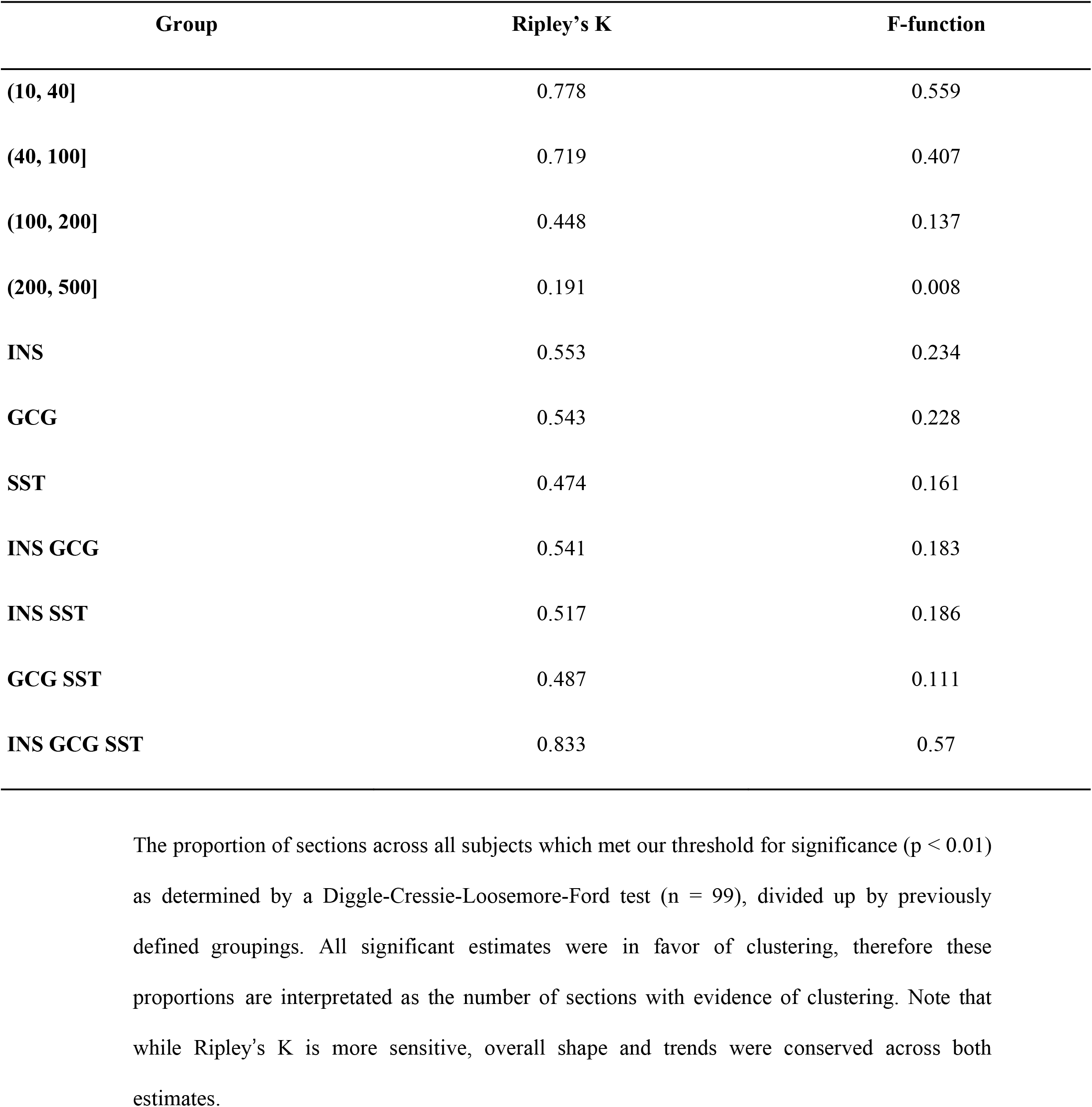
Proportion of significant values for spatial statistics.

| **Group** | **Ripley’s K** | **F-function** |
|---|---|---|
| **(10, 40]** | 0.778 | 0.559 |
| **(40, 100]** | 0.719 | 0.407 |
| **(100, 200]** | 0.448 | 0.137 |
| **(200, 500]** | 0.191 | 0.008 |
| **INS** | 0.553 | 0.234 |
| **GCG** | 0.543 | 0.228 |
| **SST** | 0.474 | 0.161 |
| **INS GCG** | 0.541 | 0.183 |
| **INS SST** | 0.517 | 0.186 |
| **GCG SST** | 0.487 | 0.111 |
| **INS GCG SST** | 0.833 | 0.57 |

Tri-hormone islets followed a similarly consistent pattern, showing significant evidence of clustering in 83.3% of sections (Table 1). Their H(r) and F(x) plots are also qualitatively similar to those derived from niSSCs and 40 – 100 μm islets; they approximate a logarithmic and fishhook shape respectively (Figure 4C). Likewise, estimates often exceeded the significance threshold from approximately 250 μm onward (Figure 4C). In contrast, estimates for the other size bins showed only sporadic and inconsistent evidence of clustering (Figure 4D, Table 1). 100 – 200 μm diameter islets clustered more frequently than 200 – 500 μm diameter islets. They achieved significance in 44.8% of sections as opposed to just 19.1% for larger islets (Table 1).

Mono-hormone subpopulations showed greater variability. Significant clustering of mono-hormone islets was observed in occasional sections throughout each pancreas (Figure 5A–C). In cases where estimates crossed the significance threshold, H(r) values were significantly elevated for all radii greater than approximately 500 – 1000 µm (Figure 5A-C). Although H(r) plots derived from these populations sometimes resembled those of tri-hormone islets, those associated with particularly strong clustering more closely approximate linear or square-root functions. Regardless of shape, actual H(r) values were consistently higher than those computed from niSSCs, 100 – 200 μm islets, or tri-hormone islets. Corresponding F(x) plots exhibit a widened fishhook shape. These data suggest intermittent but strong clustering among these subpopulations. Results were consistent across estimates of both Ripley’s K and the empty-space function (Figure 4A-D, Figure 5A-C). Estimates of F(x) were less sensitive but overall tracked with estimates of Ripley’s K (Table 1). No major discrepancies appeared between the two approaches.

**Figure 5:**
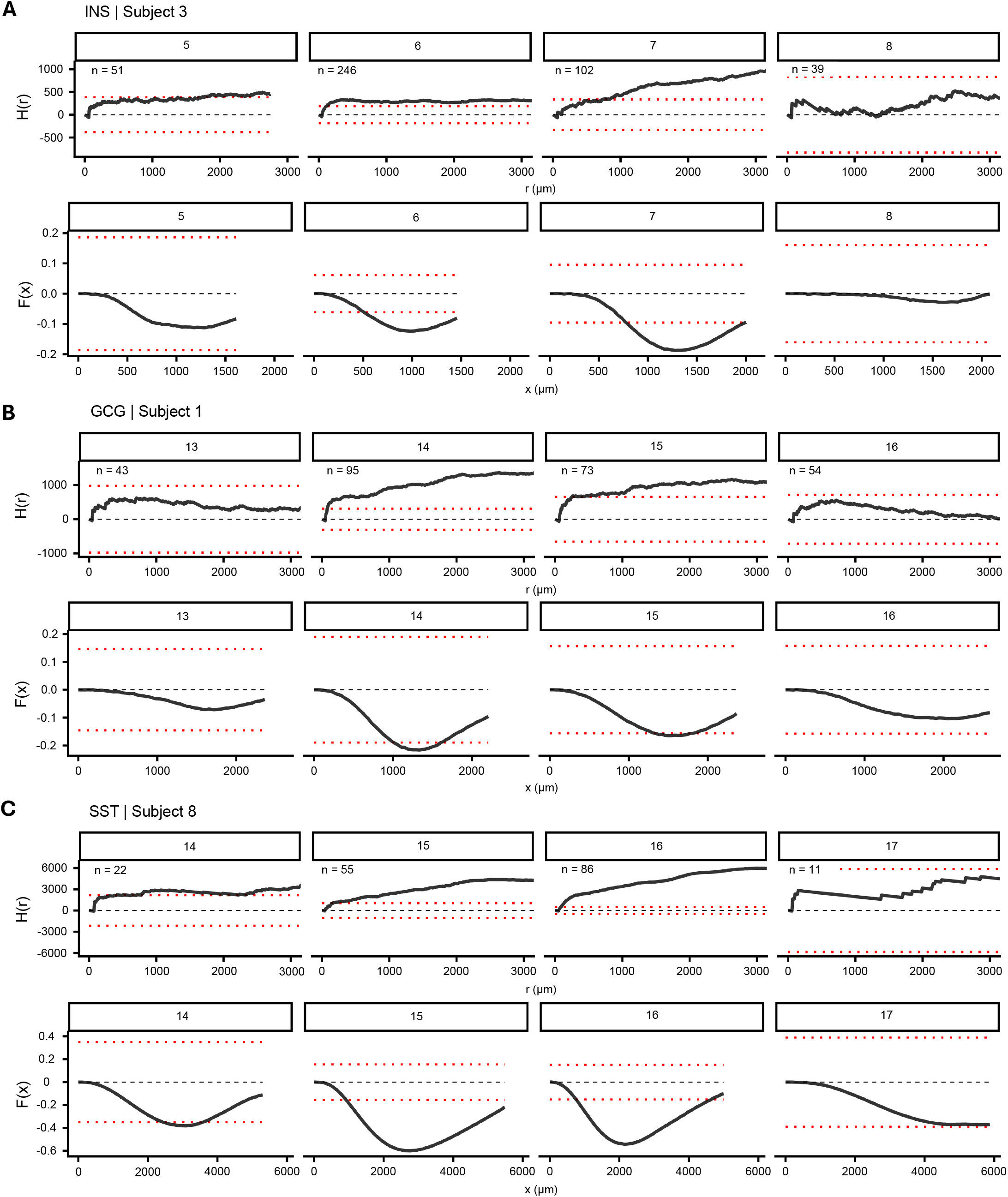
Spatial analysis of mono-hormone islets. (**A**) Representative spatial plots estimated using INS-only islets. Ripley’s K simplified to the H function is presented on the top and the empty space function F(x) is presented on the bottom. The expected value under CSR for both functions is 0. The red dotted lines show a global confidence envelope generated from 99 simulations. The header and sub-header indicate subject and section number respectively. Interpretations of the plots are reversed: An H(r) value above what is expected under CSR indicates clustering, while an F(x) value below what is expected under CSR indicates clustering. (**B**) Representative spatial plots estimated using GCG-only islets. (**C**) Representative spatial plots estimated using SST-only islets.

Significant clustering was associated with high islet counts (Figure 5A-C, Table 1). Although this association may reflect an underlying biological phenomenon, it may also result from our conservative confidence envelopes. For example, several sections that did not meet the threshold for significance in Figure 5 show patterns and values similar to those of sections that did. In these cases, the difference appears to reflect wider confidence envelopes resulting from lower islet counts rather than a different spatial distribution. Because these statistics are complex to interpret, and because of the volume of data included here, summarization necessarily obscures some underlying nuance. The full set of individual plots is therefore available upon request.

### SST-only and GCG-SST islets are found close to the border, tri-hormone islets are localized to the core

To further characterize the spatial organization of islet subpopulations, we calculated the distance from each point to the border. This computation was performed using the same representative point patterns described above. Borders in these patterns do not describe biological boundaries, rather they are derived from the points themselves (see methods for further details). Therefore, raw distance values did not have biological relevance and could be transformed without losing relevant information. Values were scaled, per-section, to fall between 0 and 1 to enable comparison between sections with different areas. Mean border distance for each hormone-based subpopulation was then quantified per-section, per-donor, and using all islets pooled together. Examination of the per-section means revealed a high degree of intra-individual variability (Figure 6A). Mean values among mono-hormone islets were particularly inconsistent and displayed large shifts from section to section. In contrast, mean values among tri-hormone islets appeared stable across sections (Figure 6A). GCG-SST islets appeared more variable than the other two di-hormone subpopulations (Figure 6A).

**Figure 6:**
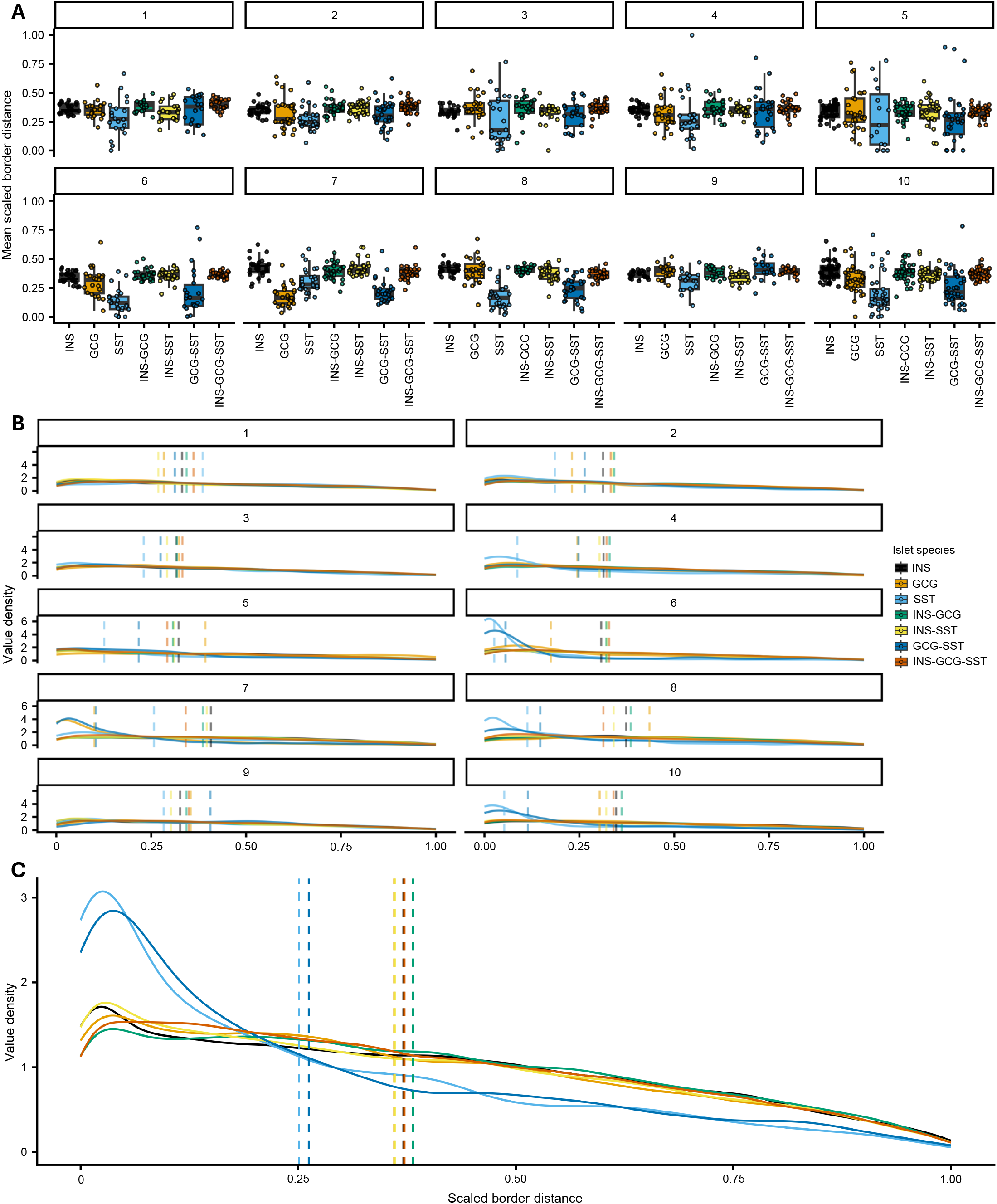
Spatial localization of mono-SST, GCG-SST and tri-hormone islets. (**A**) Mean normalized border distance per subpopulation per section. Each point represents the mean border distance in a single section. Colors denote subpopulation. Values are separated by subject, indicated by plot header. (**B**) Mean normalized border distance per subpopulation. Underlying distributions are shown via a value density function and mean distances for each subject are shown with dotted lines. (**C**) Mean normalized border distance per subpopulation with all data combined.

Zooming out to examine all islets from a donor together, inter-subject variability also became clear (Figure 6B). Both means and underlying distributions were visualized to allow for a detailed examination of the data. While the specific values and order of the subpopulation means were different between donors, the mean border distance of tri-hormone islets was never the lowest (Figure 6B). This is consistent with tri-hormone islets being located near the center of the tissue section. Underlying distributions were generally flat, though SST and/or GCG-SST islets had a very high density of low values in subjects 4, 6, 8, 7, and 10 (Figure 6B).

Lastly, data from all subjects were pooled together. This combined dataset revealed a clear distinction in means between SST and GCG-SST islets and the other subpopulations (Figure 6C). Underlying distributions likewise showed a large density of low values among these two subpopulations (Figure 6C). Comparing these results with the per-donor data suggests that these differences are driven by a select few subjects with an extremely high number of GCG and GCG-SST islets close to the border. A Kruskal-Wallis test performed on these data revealed a significant difference present between subpopulations (p < 0.01), though the effect size was small (*η_H_*^2^ = 0.0121).

## Discussion

SC-islets are manufactured from well-characterized, quality-controlled human pluripotent stem cell master and working cell banks under cGMP conditions, enabling standardized and scalable production (6). One of the critical challenges in producing therapeutic SC-islets is achieving reproducible organoid size and cellular composition, as these factors strongly influence nutrient and oxygen diffusion, cell-cell interactions and ultimately functional beta-cell maturation and glucose responsiveness (1, 2, 6, 34–36). Over the past decade, there has been a major shift in our understanding of pancreatic islet biology (21, 32). Rather than viewing beta-cells as autonomous glucose sensors, current evidence indicates that alpha- and delta-cells actively regulate beta-cell function through paracrine signaling, contributing to the establishment and maintenance of the glycemic set point (21, 32, 33, 37, 38). There is no single consensus organoid size or endocrine cell composition used to generate SC-islets. Different protocols (1, 2, 6, 39–41) produce aggregates with varied diameters and endocrine cell ratios. However, relatively consistent ranges for SC-islet size are ∼100-150 µm (34, 42) and ∼100-200 µm (43, 44). Cellular composition for both is approximately 55-70% beta-cells, 20-35% alpha-cells and 5-10% delta-cells (34, 42–44). Notably, size-dependent differences in islet cellular composition have not been accounted for. Furthermore, the consistently lower reported proportion of alpha-cells relative to beta-cells reflects the prevailing beta-cell-centric perspective and the generally accepted hierarchy of islet cell types (45–47).

Retrospectively, we reported such an unexpectedly high percentage of alpha-cells in large human islets over a decade ago (48), when only a few years later, the first seminal papers on the generation of functional SC-beta-cells/islets were out in the field (1, 2). Naturally, it was beyond our imagination at that time that such information regarding islet cellular composition would ever be meaningful in the future. Recent advances in islet physiology and biology have established the pivotal roles of intraislet paracrine signaling by alpha- and delta-cells in the regulation of insulin secretion. (49–54). As demonstrated in the current study, many islets contain an equal or even greater proportion of alpha-cells than beta-cells, with each cell type accounting for approximately 40% of the total cell mass. This comparable cellular representation underscores the close functional relationship between alpha- and beta-cells. Further, the emergence of this proportional intersection in islets larger than approximately 200 μm in diameter suggests an unexplored, size-dependent form of fine-tuning in islet composition. A consistent high ratio of delta-cells (∼17%) across the islet size distribution suggests a coordinated relationship between delta-cell abundance and the overall beta- and alpha-cell mass.

The cellular heterogeneity of islets has recently been recognized by the field (25, 27, 28). While rodent islets form a core of beta-cells accompanied by peripheral alpha- and delta-cells, there is no stereotypical adult human islet. Non-beta-cells are instead intermingled with beta-cells with various ratios. Indeed, we observe a large degree of heterogeneity in the composition of individual islets in these data. However, importantly, a majority of endocrine cells (> 80%) form islets composed of all three major islet cells. Diversity does not entirely govern the biological basics, as the islet has obviously been first designed with careful arrangement of both synergistic and antagonistic effects which require all three major islet cell types.

Even more recently, attention has been paid to mono-hormone positive cells/clusters/islets (11). While beta-cell-only islets appear to be the most common, and can most easily be linked to potential regulatory function (11), the presence of alpha-cell- and delta-cell-only islets raises equally interesting questions. Both cell types have been observed to exert regulatory effects on the exocrine pancreas (55–58). The relative abundance of these independent cells/clusters may therefore reflect some local action on the surrounding acinar cells.

Adding to this intrigue is our observation that mono-hormone islets appear to cluster together. This was true for all three cell types, and some degree of clustering in beta-cell-only islets is supported by prior work (11). There are two non-exclusive explanations for this behavior: (*i*) These cells are forming and aggregating in the mature pancreas, and/or (*ii*) these clusters are a residual consequence of early islet development. Previous investigators have proposed the aggregation model in relevance to beta-cell/islet neogenesis in particular (13), and there is some limited evidence supporting the model (59). There is likewise evidence of islet cells clustering during human development (60).

Overall, these data provide attest to the relevance of inter-islet interactions in pancreatic lobules. Within these lobules, blood flow is both closely integrated and bidirectional between the endocrine and exocrine compartments (61–63). As most pancreatic islets lack direct arteriole connections, groups of islets and neighboring acinar cells within a region may be exposed to a similar plasma environment. This could provide a potential role for mono-hormone islets in shaping and regulating these local tissue microenvironments. Such shared exposure could contribute to sensory or adaptive functions of pancreatic tissue and may help explain why endocrine and exocrine compartments are jointly affected in several pancreatic disorders, including diabetes (63).

Collectively, these findings highlight the importance of islet spatial organization in shaping coordinated pancreatic function. The spatial methods presented here provide researchers with a description and example of the tools that can be used to understand this organization. Expansion of the dataset through spatial transcriptomics and proteomics may also assist in forming a more complete picture of the microenvironments present in the pancreas.

## Methods

### Sex as a biological variable

Our study exclusively examined male humans. It is unknown whether the findings are relevant for female humans.

### Human pancreas specimens

Human pancreata from donors were provided by the Gift of Hope Organ Procurement Organization (Chicago, Illinois, USA).

### Pancreatic tissue imaging

Ten non-diabetic male subjects (age: 38-48 years old and BMI: 16.4 to 34.8) were studied (Supplementary Table 1). All but one of these subjects (subject 7) have been previously characterized and published (10, 25). Original images and scripts are available in those works (10, 25, 64). Each pancreas was divided into consecutive tissue blocks (∼5 mm) with alternating collection between paraffin-embedding and snap frozen. In this study, paraffin-embedded sections (5 μm) were stained with the following primary antibodies (all 1:500): polyclonal guinea pig anti-porcine insulin ([RRID]: AB_964170; Novus Biologicals), mouse monoclonal anti-human glucagon (AB_259852; MilliporeSigma), polyclonal goat anti-somatostatin (AB_2915922; Everest Biotech), and DAPI (AB_2869624; BD Biosciences). The primary antibodies were detected using a combination of DyLight 488, 549 and 649-conjugated secondary antibodies (1:200, Jackson ImmunoResearch Laboratory, West Grove, PA). Microscopic images were taken with an Olympus IX8 DSU spinning disk confocal microscope (Melville, NY) with imaging software StereoInvestigator (MicroBrightField, Williston, VT). Image processing was performed with custom scripts written for ImageJ/Fiji. This script performed automated contouring of islet areas after thresholding to include unstained fractions such as vascular structures. Each islet was given an ID and the centroid coordinates, area, and individual hormone areas were calculated. A video of the process and the original scripts are available (10, 25, 64).

### Data preprocessing

Data obtained from the automated imaging pipeline were processed and tagged for region and hormone positivity. Each islet’s effective diameter was computed based on its area. Only islets with IEDs between 10 and 500 μm were considered in our final dataset. An islet was marked as positive for a hormone only if the fluorescent area for the respective marker was greater than 40 μm^2^ to avoid inclusion based on visual artifacts. Islets which did not meet these thresholds for any of the three hormones were excluded from consideration. The first 20% of the tissue blocks were designated as the head, with the remaining 80% split in half to form the body and tail. Preference was given to the tail in the case of an uneven split. Finally, islets were binned by diameter. Two different sets of bins were used: A finer set, based on a logarithmic scale of diameter divided by the diameter of a single cell (64), and a coarser set intended to sort islets into clusters, small islets, medium islets, and large islets. Endocrine cells with diameter of 10-40 μm were additionally marked as niSSCs. The final data table included total islet area, fluorescent area, X-Y coordinates of each islet centroid, subject ID, region, original tissue block number, bin, hormone positivity, and INS, GCG, and SST fluorescent areas.

### Pancreas Characterization

All statistics presented were calculated using base R (65) along with ggplot2 (66) for presentation and tidyverse (67) for data manipulation. The packages spatstat (68), sf (69), and ggforce (70) were also used for spatial calculations. ggpubr (71), ggbeeswarm (72), and cowplot (73) were used for aesthetics and figure creation. All per-islet calculations (i.e. average fraction of INS-positive area in an islet) were computed with total fluorescent area as the denominator, not the total area of the islet structure. This statistic serves as a measurement of the composition of endocrine cells specifically and does not include area occupied by vasculature or lipid pockets.

### Spatial statistics

Ripley’s K was used to assess the spatial organization of islets within a single section of the pancreas. This function computes the number of extra events found in circles of increasing radii centered on each point as compared to what would be expected based purely on the event (islet) density (74). To assess the likelihood that a given estimate is significant, empirical values are compared to the results of 99 simulations of Poisson processes in the same plane with the same number of points. These simulated patterns approximate complete spatial randomness (CSR) to generate a 99% confidence interval for the null hypothesis of random dispersion between islets (74, 75). A concave hull was fit to the points which approximated the tissue border using the concaveman method in the sf package. Values for r were computed per-section by dividing the maximum dimension of a rectangle bounding the tissue hull by 4 (75), then creating a sequence of 513 radii from 0 to that value. In the interest of interpretability, a series of transformations were also applied. First, the output was converted to the linear function L(r) using spatstat’s inbuilt Lest function. Then, r was subtracted from each respective L(r) output to make the null value approximately 0 (75). The final presented data are in this form, known as the H function or H(r). The empty-space function was also used to assess spatial organization within a single slice. This function estimates the distribution of the distance between a random, non-event point on the plane and an event (76). The value is estimated by measuring the distance to the nearest event from n random locations in the plane. We again ran this function repeatedly on simulations of CSR to generate a 99% confidence interval. The final values here are presented as a cdf plot.

### Border distance

Border distance was computed using subpopulations of islets. For these estimates, the same tissue hull – fitted via concaveman – was used. This means that the raw distances obtained do not have direct biological significance and can comfortably be transformed without a loss of information. We therefore standardized the values obtained for each section to fall between 0 and 1.

### Statistics

Data are presented as means ± one standard deviation (SD), median is presented ± mean absolute deviation. One-way ANOVAs were performed across each region’s hormone density and across the proportions of total hormone area in each region. A simple linear regression was applied to islets using IED as the predictor for INS-, GCG-, and SST-fractions respectively. Confidence intervals for Ripley’s K and the empty-space function are described above. A Diggle-Cressie-Loosemore-Ford test was performed with 99 simulations to test for deviations from CSR. This value was chosen due to the computational cost per simulation. In order to assess the significance of border-group differences, we performed a Kruskal-Wallis test on our observed data and on 999 permutations per section. The values of the test statistic in each permutation were ranked against the empirical value and used to derive a p-value. Effect size was computed via ordinal eta-squared (Eq. 1),

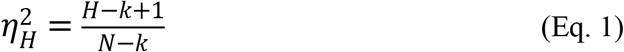

where H refers to the test statistic, k is the number of groups, and N is the number of observations (77). A p-value at or less than 0.01 was considered significant.

### Study approval

The use of deidentified human tissues in the study was approved by the IRB of the University of Chicago. For the use of human pancreas samples in this study, written informed consent was obtained from the donor or their next of kin. The specimens were all deidentified.

### Data availability

All scripts used can be found in the lab’s repository (https://github.com/Hara-Lab201/Sean). Values for all data points in graphs are reported in the Supporting Data Values file.

## Author contributions

MH conceived the idea and designed the study. SDH and MH performed experiments. SDH and VP analyzed data. SDH and MH wrote the manuscript. MH is the guarantor of this work and, as such, had full access to all the data in the study and takes responsibility for the integrity of the data and the accuracy of the data analysis.

## AI disclosure

The authors declare that no AI tools were used in the original research for or writing of this article.

## Acknowledgments

This research was supported (VP) by the Intramural Research Program of the National Institute of Diabetes and Digestive and Kidney Diseases (NIDDK) within the National Institutes of Health (NIH). The contributions of the NIH author(s) are considered Works of the United States Government. The findings and conclusions presented in this paper are those of the author(s) and do not necessarily reflect the views of the NIH or the U.S. Department of Health and Human Services. We thank Piotr Witkowski, J. Michael Millis, and Martin Jendrisak of Gift of Hope Organ Procurement Organization (Chicago, Illinois, USA) for providing the human pancreas Tissues.

