## Supplemental Figures for "Islet size, cellular composition, and spatial location are interconnected in the human pancreas"

### Slide 1
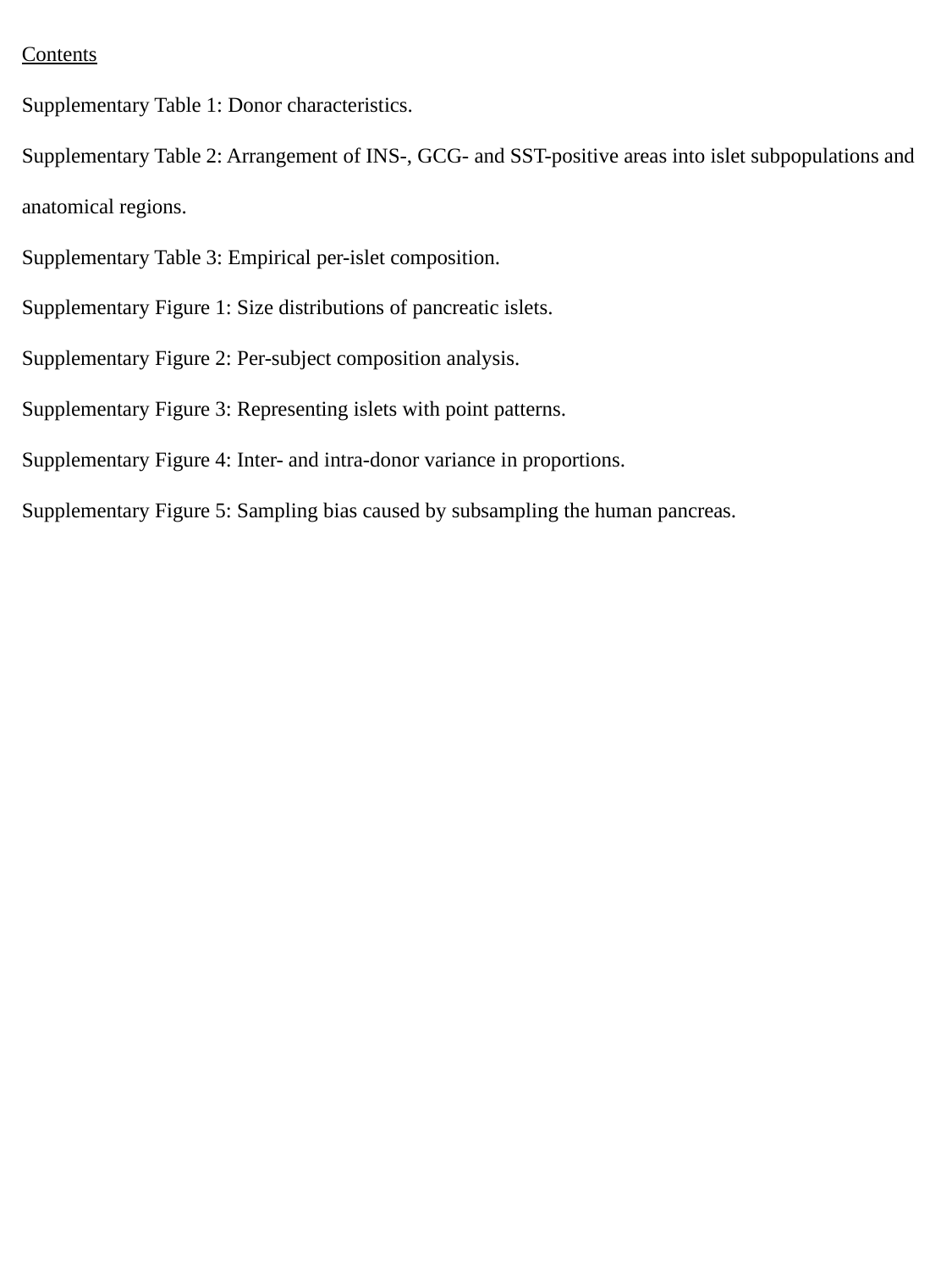

Contents
Supplementary Table 1: Donor characteristics.
Supplementary Table 2: Arrangement of INS-, GCG- and SST-positive areas into islet subpopulations and anatomical regions.
Supplementary Table 3: Empirical per-islet composition.
Supplementary Figure 1: Size distributions of pancreatic islets.
Supplementary Figure 2: Per-subject composition analysis.
Supplementary Figure 3: Representing islets with point patterns.
Supplementary Figure 4: Inter- and intra-donor variance in proportions.
Supplementary Figure 5: Sampling bias caused by subsampling the human pancreas.

### Slide 2
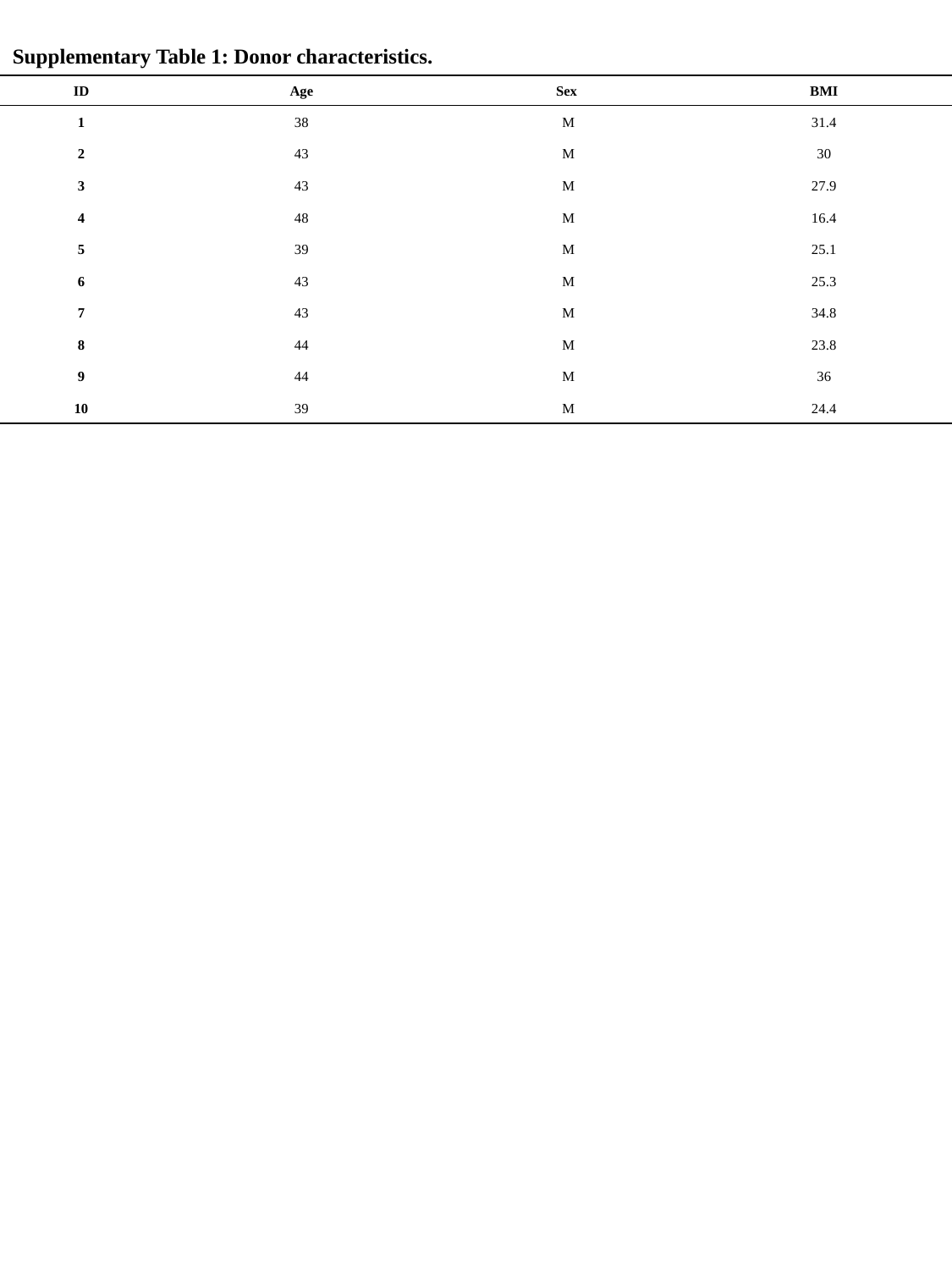

Supplementary Table 1: Donor characteristics.
| ID | Age | Sex | BMI |
| --- | --- | --- | --- |
| 1 | 38 | M | 31.4 |
| 2 | 43 | M | 30 |
| 3 | 43 | M | 27.9 |
| 4 | 48 | M | 16.4 |
| 5 | 39 | M | 25.1 |
| 6 | 43 | M | 25.3 |
| 7 | 43 | M | 34.8 |
| 8 | 44 | M | 23.8 |
| 9 | 44 | M | 36 |
| 10 | 39 | M | 24.4 |

### Slide 3
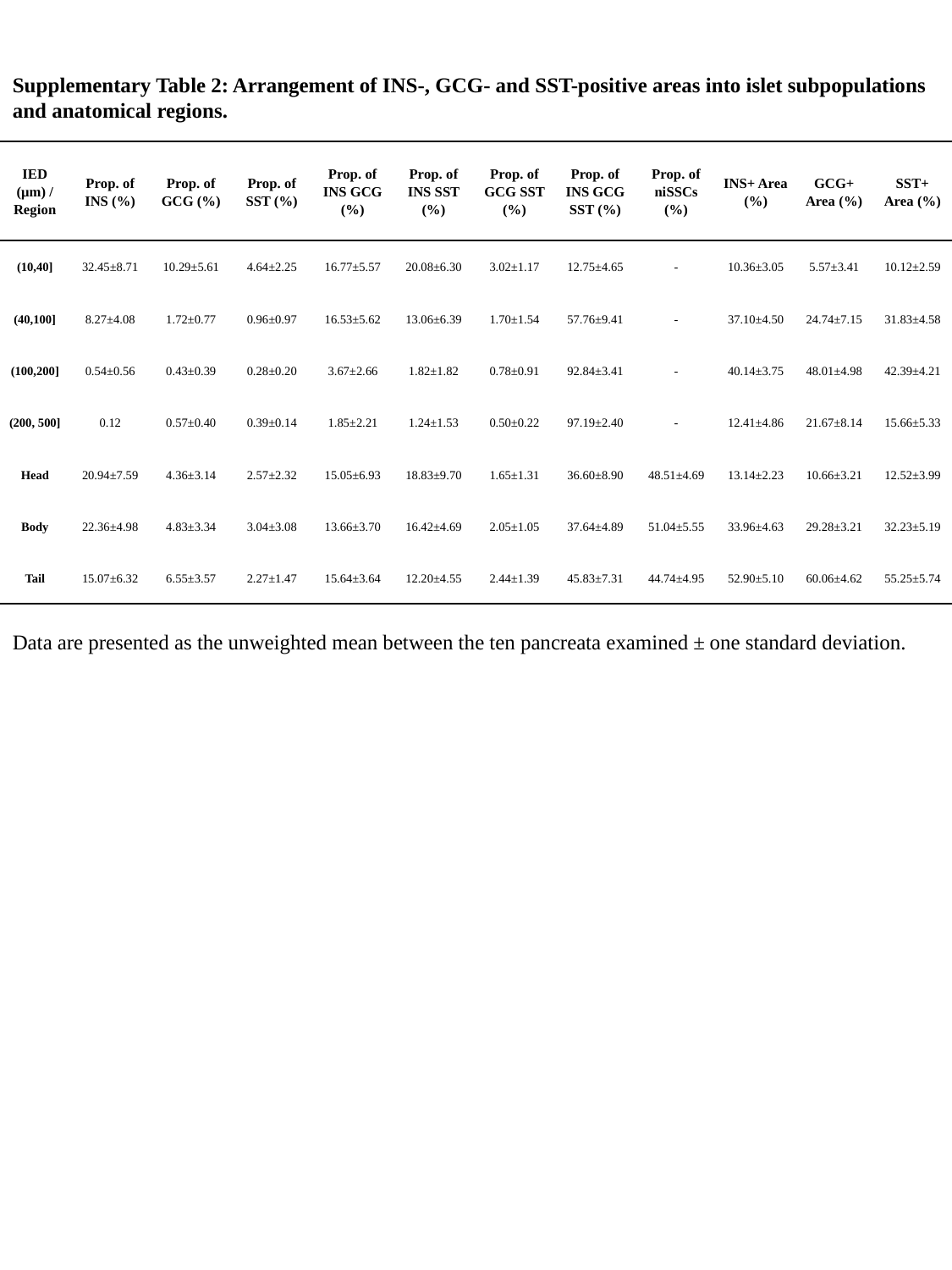

Supplementary Table 2: Arrangement of INS-, GCG- and SST-positive areas into islet subpopulations and anatomical regions.
| IED (μm) / Region | Prop. of INS (%) | Prop. of GCG (%) | Prop. of SST (%) | Prop. of INS GCG (%) | Prop. of INS SST (%) | Prop. of GCG SST (%) | Prop. of INS GCG SST (%) | Prop. of niSSCs (%) | INS+ Area (%) | GCG+ Area (%) | SST+ Area (%) |
| --- | --- | --- | --- | --- | --- | --- | --- | --- | --- | --- | --- |
| (10,40] | 32.45±8.71 | 10.29±5.61 | 4.64±2.25 | 16.77±5.57 | 20.08±6.30 | 3.02±1.17 | 12.75±4.65 | - | 10.36±3.05 | 5.57±3.41 | 10.12±2.59 |
| (40,100] | 8.27±4.08 | 1.72±0.77 | 0.96±0.97 | 16.53±5.62 | 13.06±6.39 | 1.70±1.54 | 57.76±9.41 | - | 37.10±4.50 | 24.74±7.15 | 31.83±4.58 |
| (100,200] | 0.54±0.56 | 0.43±0.39 | 0.28±0.20 | 3.67±2.66 | 1.82±1.82 | 0.78±0.91 | 92.84±3.41 | - | 40.14±3.75 | 48.01±4.98 | 42.39±4.21 |
| (200, 500] | 0.12 | 0.57±0.40 | 0.39±0.14 | 1.85±2.21 | 1.24±1.53 | 0.50±0.22 | 97.19±2.40 | - | 12.41±4.86 | 21.67±8.14 | 15.66±5.33 |
| Head | 20.94±7.59 | 4.36±3.14 | 2.57±2.32 | 15.05±6.93 | 18.83±9.70 | 1.65±1.31 | 36.60±8.90 | 48.51±4.69 | 13.14±2.23 | 10.66±3.21 | 12.52±3.99 |
| Body | 22.36±4.98 | 4.83±3.34 | 3.04±3.08 | 13.66±3.70 | 16.42±4.69 | 2.05±1.05 | 37.64±4.89 | 51.04±5.55 | 33.96±4.63 | 29.28±3.21 | 32.23±5.19 |
| Tail | 15.07±6.32 | 6.55±3.57 | 2.27±1.47 | 15.64±3.64 | 12.20±4.55 | 2.44±1.39 | 45.83±7.31 | 44.74±4.95 | 52.90±5.10 | 60.06±4.62 | 55.25±5.74 |
Data are presented as the unweighted mean between the ten pancreata examined ± one standard deviation.

### Slide 4
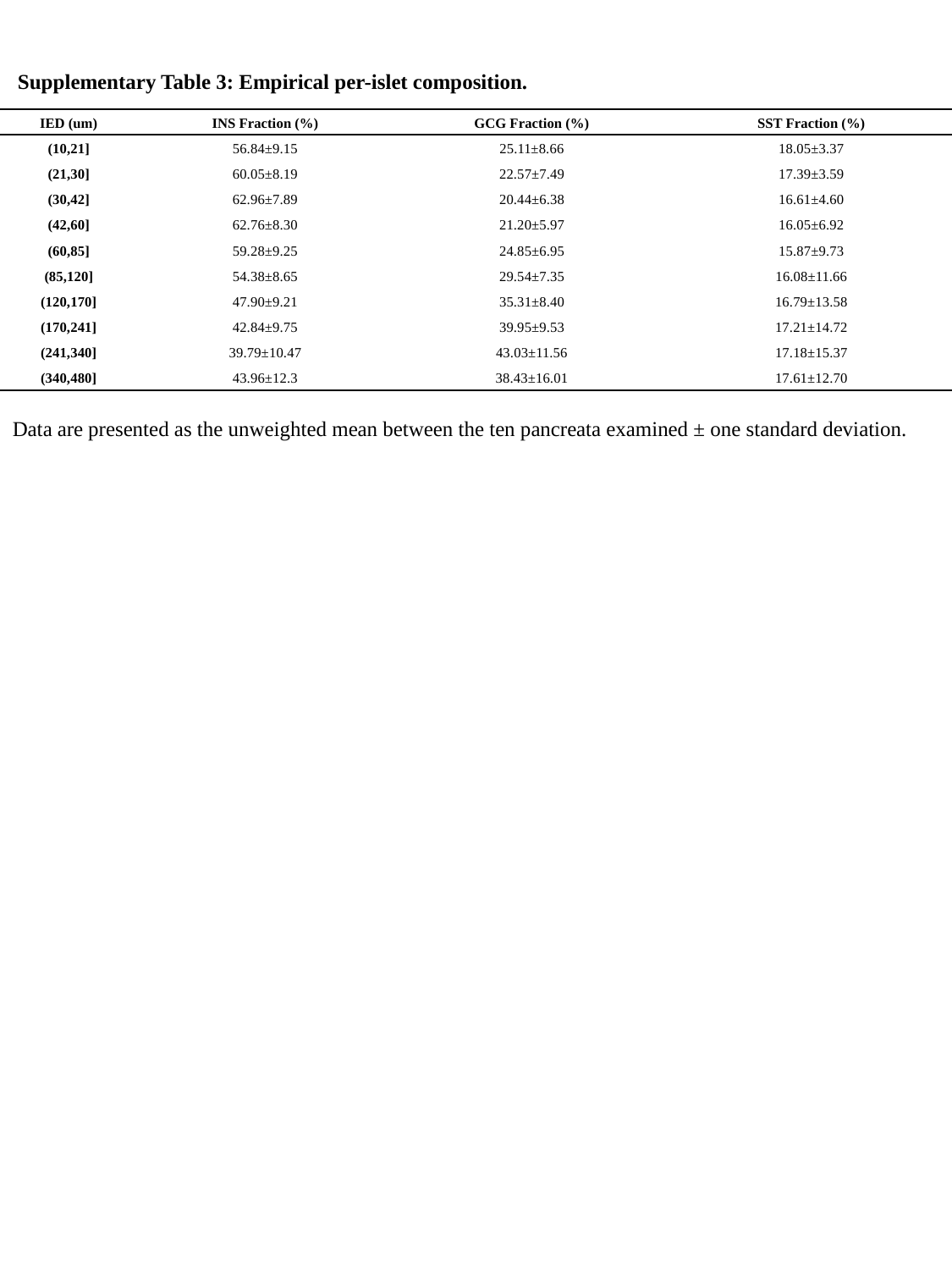

Supplementary Table 3: Empirical per-islet composition.
| IED (um) | INS Fraction (%) | GCG Fraction (%) | SST Fraction (%) |
| --- | --- | --- | --- |
| (10,21] | 56.84±9.15 | 25.11±8.66 | 18.05±3.37 |
| (21,30] | 60.05±8.19 | 22.57±7.49 | 17.39±3.59 |
| (30,42] | 62.96±7.89 | 20.44±6.38 | 16.61±4.60 |
| (42,60] | 62.76±8.30 | 21.20±5.97 | 16.05±6.92 |
| (60,85] | 59.28±9.25 | 24.85±6.95 | 15.87±9.73 |
| (85,120] | 54.38±8.65 | 29.54±7.35 | 16.08±11.66 |
| (120,170] | 47.90±9.21 | 35.31±8.40 | 16.79±13.58 |
| (170,241] | 42.84±9.75 | 39.95±9.53 | 17.21±14.72 |
| (241,340] | 39.79±10.47 | 43.03±11.56 | 17.18±15.37 |
| (340,480] | 43.96±12.3 | 38.43±16.01 | 17.61±12.70 |
Data are presented as the unweighted mean between the ten pancreata examined ± one standard deviation.

### Slide 5
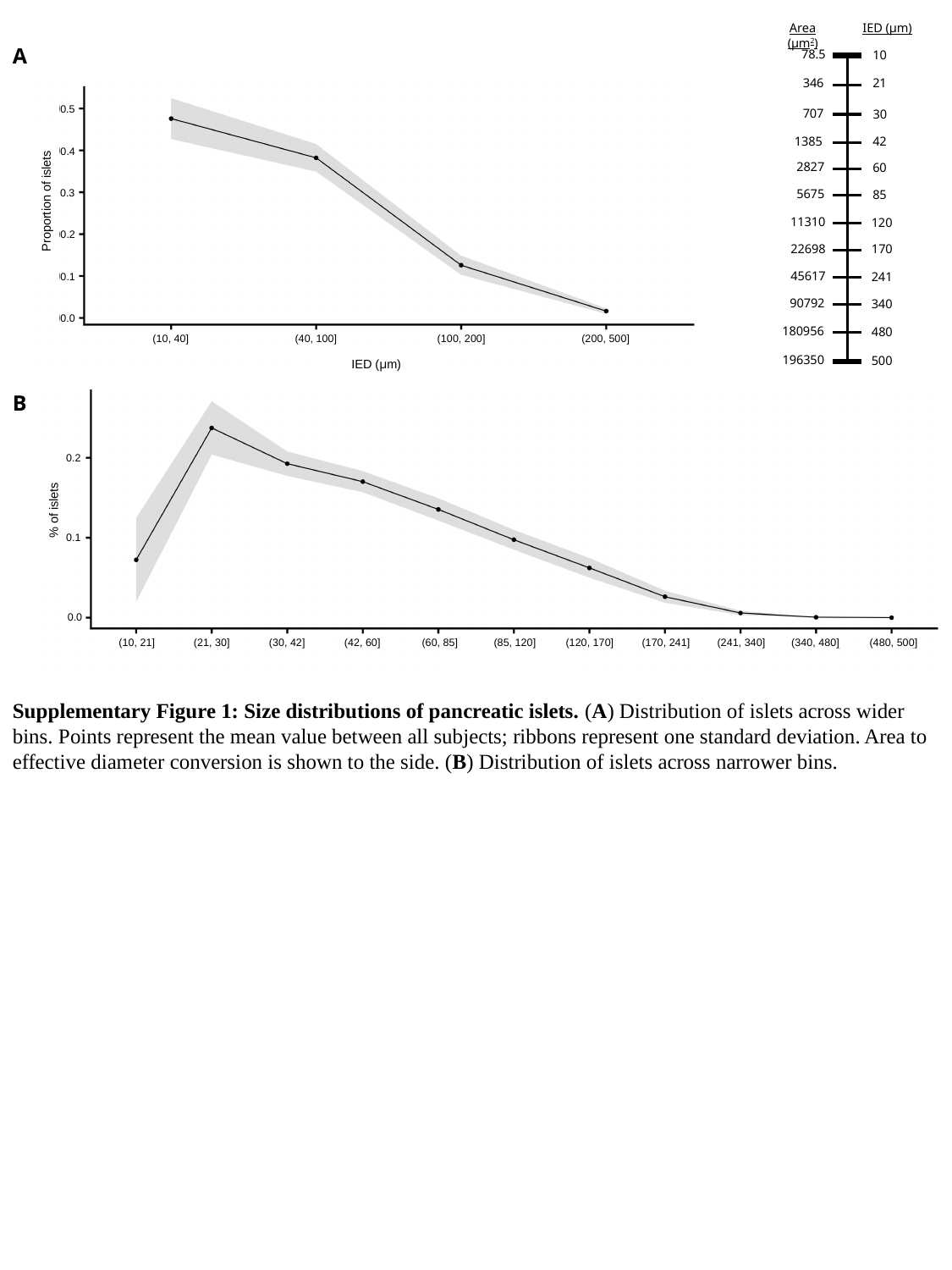

Area (μm^2^)
IED (μm)
A
0.5
500
480
340
120
241
170
30
60
42
85
21
10
707
346
78.5
2827
5675
1385
45617
22698
11310
90792
180956
196350
0.4
0.3
Proportion of islets
0.2
0.1
0.0
(100, 200]
(200, 500]
(10, 40]
(40, 100]
IED (μm)
B
0.2
% of islets
0.1
0.0
(480, 500]
(340, 480]
(170, 241]
(120, 170]
(85, 120]
(30, 42]
(42, 60]
(21, 30]
(10, 21]
(241, 340]
(60, 85]
Supplementary Figure 1: Size distributions of pancreatic islets. (A) Distribution of islets across wider bins. Points represent the mean value between all subjects; ribbons represent one standard deviation. Area to effective diameter conversion is shown to the side. (B) Distribution of islets across narrower bins.

### Slide 6
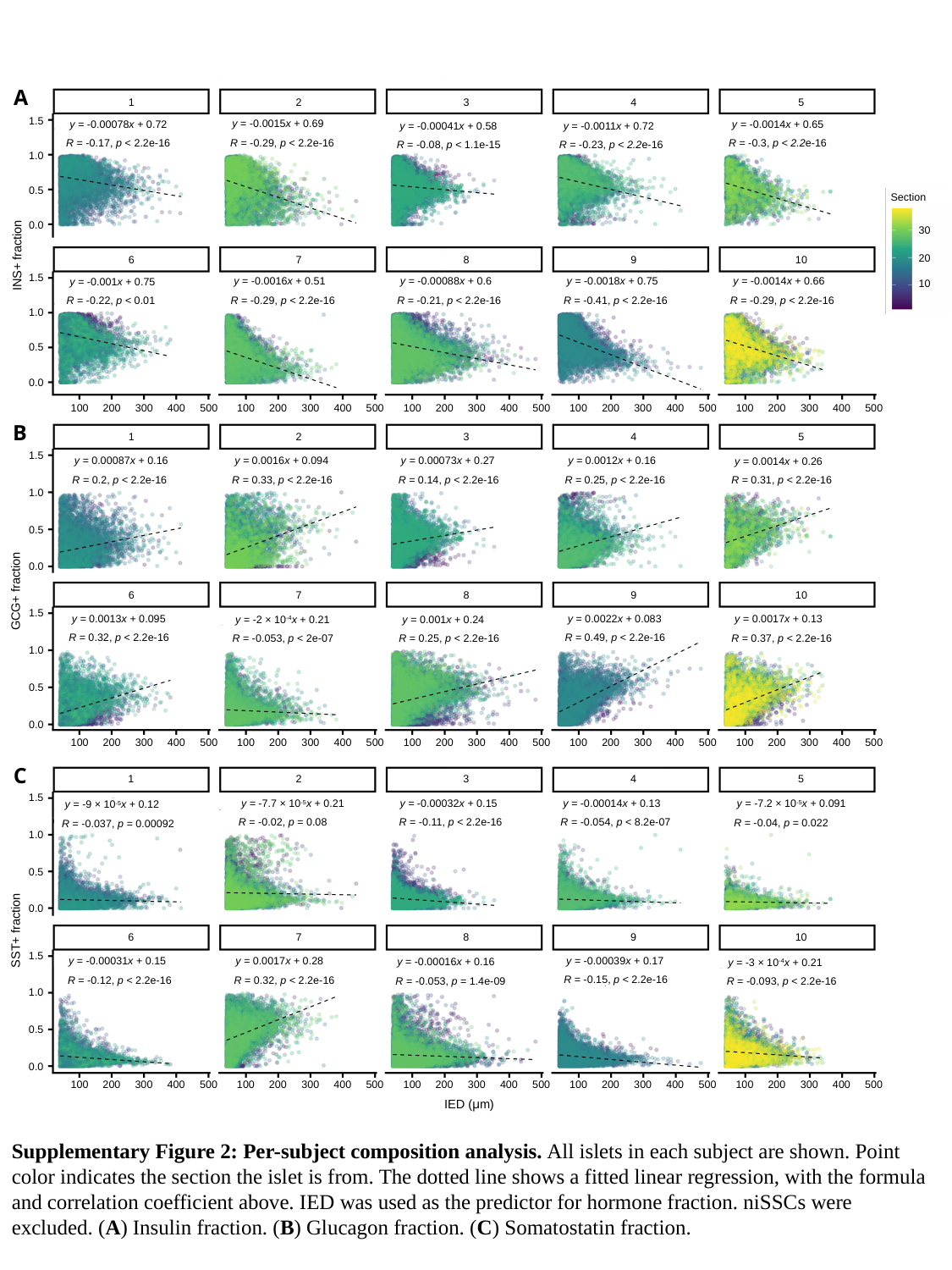

A
2
3
4
5
1
1.5
y = -0.0015x + 0.69
y = -0.00078x + 0.72
y = -0.0014x + 0.65
y = -0.0011x + 0.72
y = -0.00041x + 0.58
R = -0.29, p < 2.2e-16
R = -0.17, p < 2.2e-16
R = -0.3, p < 2.2e-16
R = -0.23, p < 2.2e-16
R = -0.08, p < 1.1e-15
1.0
0.5
Section
0.0
30
INS+ fraction
20
7
8
9
10
6
1.5
y = -0.0016x + 0.51
y = -0.00088x + 0.6
y = -0.0018x + 0.75
y = -0.0014x + 0.66
y = -0.001x + 0.75
10
R = -0.21, p < 2.2e-16
R = -0.41, p < 2.2e-16
R = -0.29, p < 2.2e-16
R = -0.29, p < 2.2e-16
R = -0.22, p < 0.01
1.0
0.5
0.0
400
500
100
300
400
500
400
500
400
500
400
500
200
100
300
100
300
100
300
100
300
200
200
200
200
B
2
3
4
5
1
1.5
y = 0.0012x + 0.16
y = 0.0016x + 0.094
y = 0.00087x + 0.16
y = 0.00073x + 0.27
y = 0.0014x + 0.26
R = 0.2, p < 2.2e-16
R = 0.25, p < 2.2e-16
R = 0.33, p < 2.2e-16
R = 0.14, p < 2.2e-16
R = 0.31, p < 2.2e-16
1.0
0.5
0.0
GCG+ fraction
7
8
9
10
6
1.5
y = 0.0017x + 0.13
y = 0.0022x + 0.083
y = 0.0013x + 0.095
y = 0.001x + 0.24
y = -2 × 10^-4^x + 0.21
R = 0.49, p < 2.2e-16
R = 0.32, p < 2.2e-16
R = 0.37, p < 2.2e-16
R = 0.25, p < 2.2e-16
R = -0.053, p < 2e-07
1.0
0.5
0.0
400
500
100
300
400
500
400
500
400
500
400
500
200
100
300
100
300
100
300
100
300
200
200
200
200
C
2
3
4
5
1
1.5
y = -0.00014x + 0.13
y = -0.00032x + 0.15
y = -7.7 × 10^-5^x + 0.21
y = -7.2 × 10^-5^x + 0.091
y = -9 × 10^-5^x + 0.12
R = -0.054, p < 8.2e-07
R = -0.11, p < 2.2e-16
R = -0.02, p = 0.08
R = -0.04, p = 0.022
R = -0.037, p = 0.00092
1.0
0.5
0.0
SST+ fraction
7
8
9
10
6
1.5
y = 0.0017x + 0.28
y = -0.00039x + 0.17
y = -0.00031x + 0.15
y = -0.00016x + 0.16
y = -3 × 10^-4^x + 0.21
R = -0.15, p < 2.2e-16
R = -0.12, p < 2.2e-16
R = 0.32, p < 2.2e-16
R = -0.053, p = 1.4e-09
R = -0.093, p < 2.2e-16
1.0
0.5
0.0
400
500
100
300
400
500
400
500
400
500
400
500
200
100
300
100
300
100
300
100
300
200
200
200
200
IED (μm)
Supplementary Figure 2: Per-subject composition analysis. All islets in each subject are shown. Point color indicates the section the islet is from. The dotted line shows a fitted linear regression, with the formula and correlation coefficient above. IED was used as the predictor for hormone fraction. niSSCs were excluded. (A) Insulin fraction. (B) Glucagon fraction. (C) Somatostatin fraction.

### Slide 7
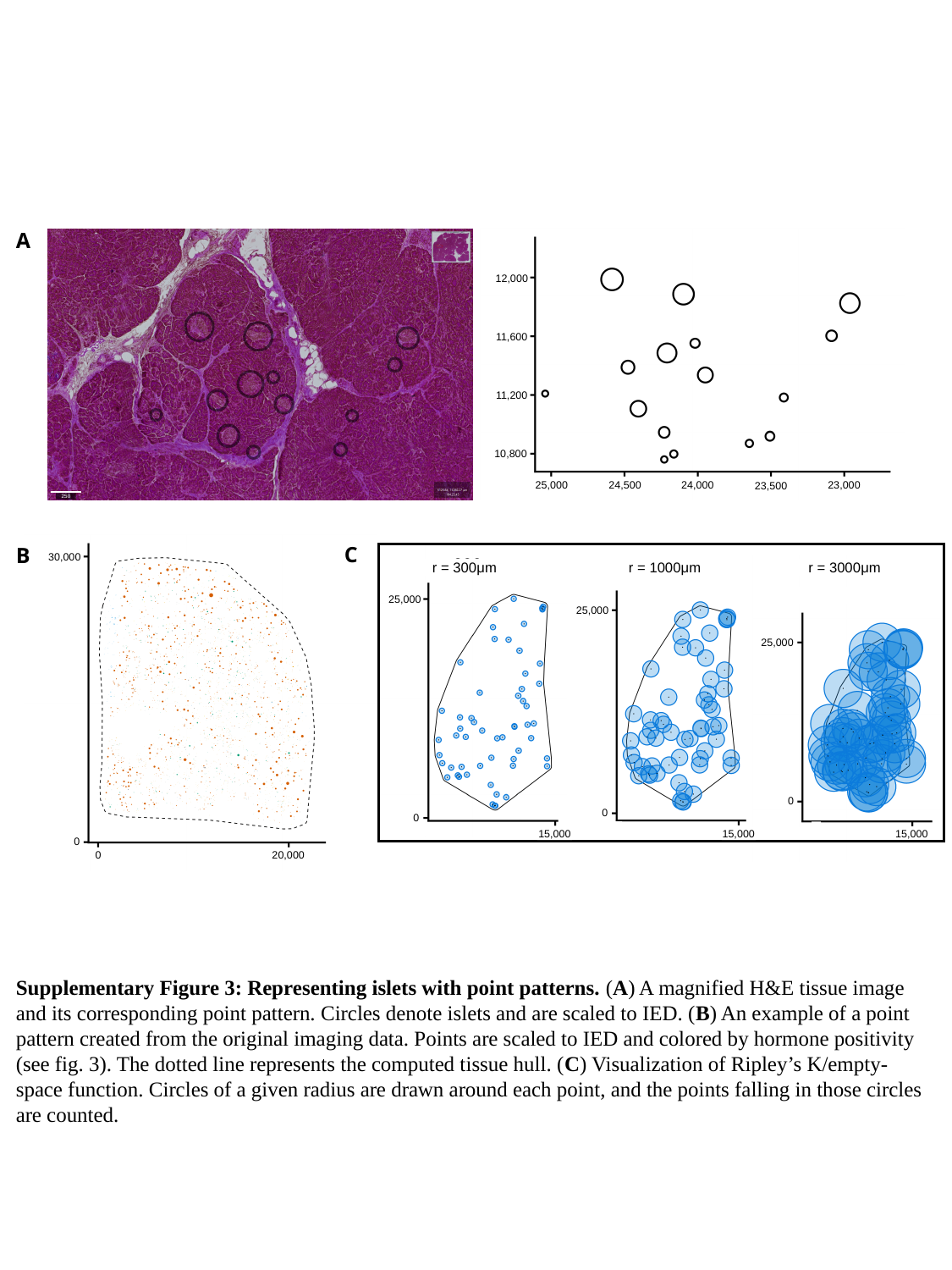

A
12,000
11,600
11,200
10,800
25,000
24,500
24,000
23,000
23,500
250
C
B
30,000
r = 300μm
r = 1000μm
r = 3000μm
25,000
25,000
25,000
0
0
0
15,000
15,000
15,000
0
0
20,000
Supplementary Figure 3: Representing islets with point patterns. (A) A magnified H&E tissue image and its corresponding point pattern. Circles denote islets and are scaled to IED. (B) An example of a point pattern created from the original imaging data. Points are scaled to IED and colored by hormone positivity (see fig. 3). The dotted line represents the computed tissue hull. (C) Visualization of Ripley’s K/empty-space function. Circles of a given radius are drawn around each point, and the points falling in those circles are counted.

### Slide 8
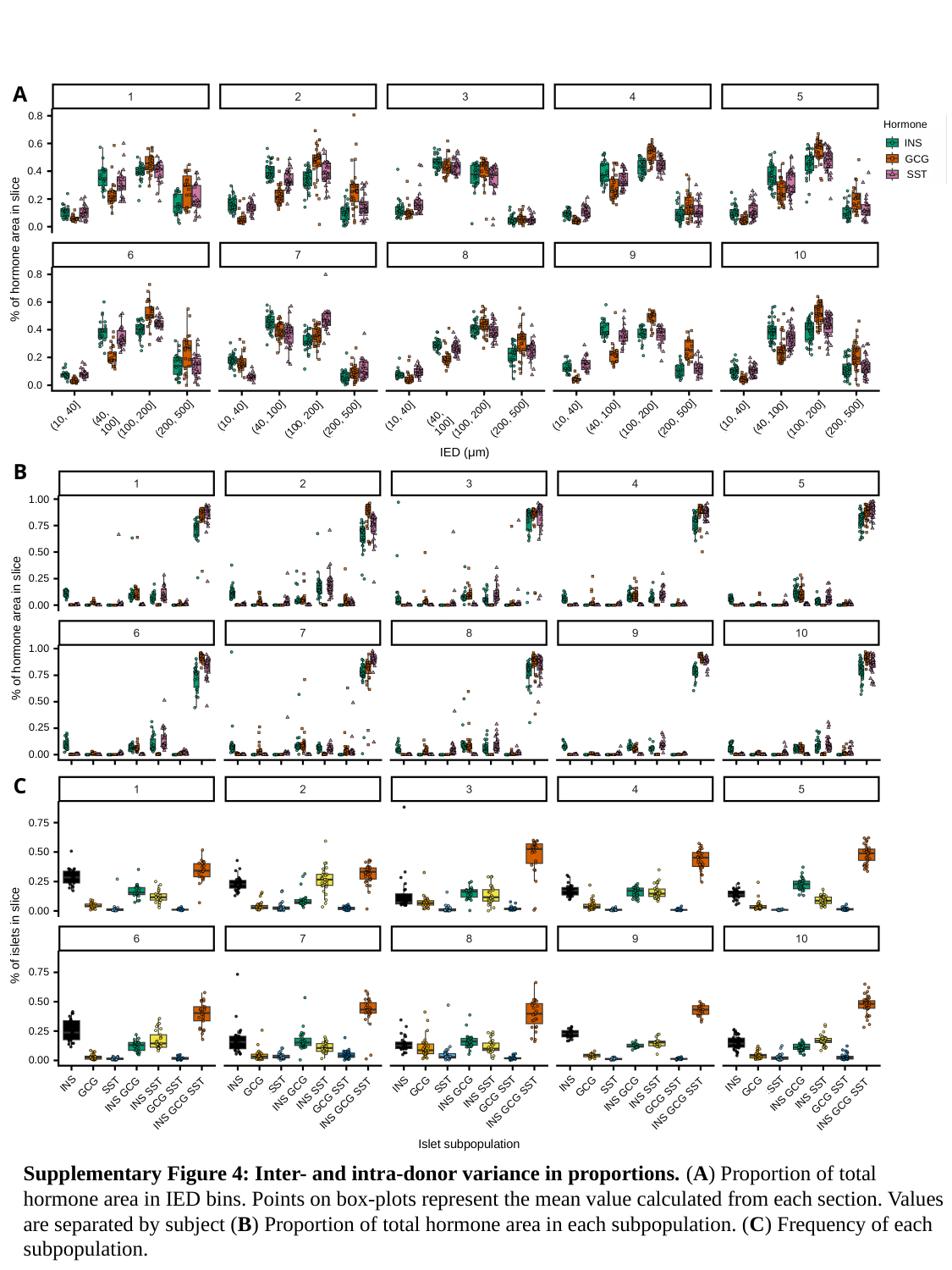

A
0.8
Hormone
INS
GCG
SST
0.6
0.4
0.2
0.0
% of hormone area in slice
0.8
0.6
0.4
0.2
0.0
(10, 40]
(10, 40]
(10, 40]
(10, 40]
(10, 40]
(40, 100]
(40, 100]
(40, 100]
(40, 100]
(40, 100]
(200, 500]
(200, 500]
(200, 500]
(200, 500]
(200, 500]
(100, 200]
(100, 200]
(100, 200]
(100, 200]
(100, 200]
IED (μm)
B
1.00
0.75
0.50
0.25
0.00
% of hormone area in slice
1.00
0.75
0.50
0.25
0.00
C
0.75
0.50
0.25
0.00
% of islets in slice
0.75
0.50
0.25
0.00
INS
INS
INS
INS
INS
SST
SST
SST
SST
SST
GCG
GCG
GCG
GCG
GCG
INS SST
INS SST
INS SST
INS SST
INS GCG
INS GCG
INS SST
INS GCG
INS GCG
GCG SST
INS GCG
GCG SST
GCG SST
GCG SST
GCG SST
INS GCG SST
INS GCG SST
INS GCG SST
INS GCG SST
INS GCG SST
Islet subpopulation
Supplementary Figure 4: Inter- and intra-donor variance in proportions. (A) Proportion of total hormone area in IED bins. Points on box-plots represent the mean value calculated from each section. Values are separated by subject (B) Proportion of total hormone area in each subpopulation. (C) Frequency of each subpopulation.

### Slide 9
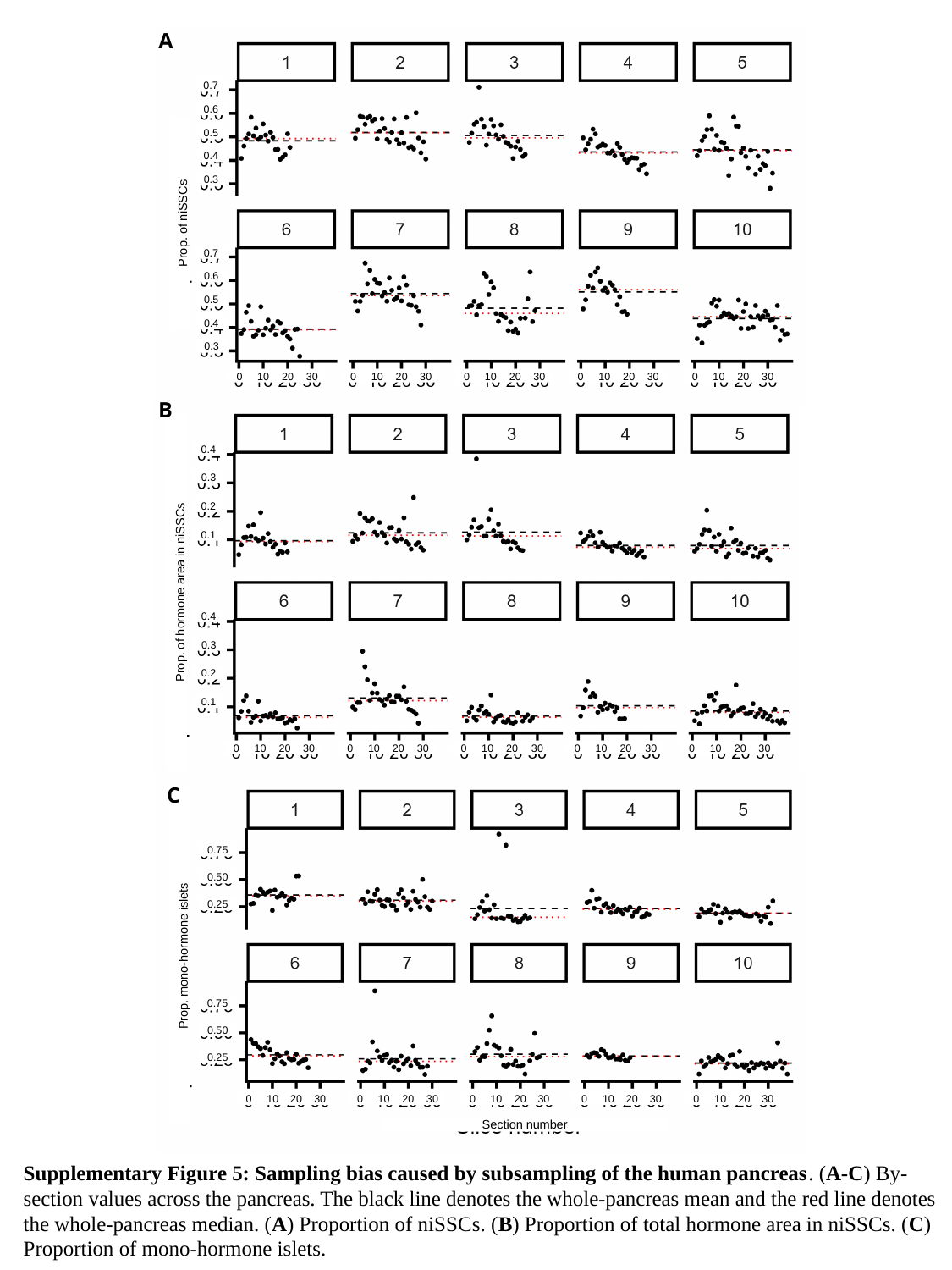

A
0.7
0.6
0.5
0.4
0.3
Prop. of niSSCs
0.7
0.6
0.5
0.4
0.3
0
10
20
30
0
10
20
30
0
10
20
30
0
10
20
30
0
10
20
30
B
0.4
0.3
0.2
0.1
Prop. of hormone area in niSSCs
0.4
0.3
0.2
0.1
0
10
20
30
0
10
20
30
0
10
20
30
0
10
20
30
0
10
20
30
C
0.75
0.50
0.25
Prop. mono-hormone islets
0.75
0.50
0.25
20
30
20
30
20
30
20
30
20
30
0
10
0
10
0
10
0
10
0
10
Section number
Supplementary Figure 5: Sampling bias caused by subsampling of the human pancreas. (A-C) By-section values across the pancreas. The black line denotes the whole-pancreas mean and the red line denotes the whole-pancreas median. (A) Proportion of niSSCs. (B) Proportion of total hormone area in niSSCs. (C) Proportion of mono-hormone islets.
